# Sets of Co-regulated Serum Lipids are Associated with Alzheimer Disease Pathophysiology

**DOI:** 10.1101/550723

**Authors:** Dinesh Kumar Barupal, Rebecca Bailli, Sili Fan, Andrew J. Saykin, Peter J. Meikle, Matthias Arnold, Kwangsik Nho, Oliver Fiehn, Rima Kaddurah-Daouk, for the Alzheimer’s Disease Neuroimaging Initiative, the Alzheimer Disease Metabolomics Consortium

## Abstract

**INTRODUCTION:** Altered regulation of lipid metabolism in Alzheimer disease (AD) can be characterized using lipidomic profiling.

**METHOD:** 349 serum lipids were measured in 806 participants enrolled in the Alzheimer Disease Neuroimaging Initiative Phase 1 (ADNI1) cohort and analysed using lipid regression models and lipid set enrichment statistics.

**RESULTS:** AD diagnosis was associated with 7 of 28 lipid sets of which four also correlated with cognitive decline, including polyunsaturated fatty acids. CSF amyloid beta Aβ_1-42_ correlated with glucosylceramides, lysophosphatidyl cholines and unsaturated triacylglycerides; CSF total tau and brain atrophy correlated with monounsaturated sphingomyelins and ceramides, in addition to EPA-containing lipids.

**DISCUSSION:** Lipid desaturation, elongation and acyl chain remodeling are dysregulated across the spectrum of AD pathogenesis. Monounsaturated lipids were important in early stages of AD, while polyunsaturated lipid metabolism was associated with later stages of AD.

**SIGNFICANCE:** Both metabolic genes and co-morbidity with metabolic diseases indicate that lipid metabolism is critical in the etiology of Alzheimer’s disease (AD). For 800 subjects, we found that sets of blood lipids were associated with current AD-biomarkers and with AD clinical symptoms. Our study highlights the role of disturbed acyl chain lipid remodelling in several lipid classes. Our work has significant implications on finding a cure for AD. Depending on subject age, human blood lipids may have different effects on AD development. Remodelling of acyl chains needs to be studied in relation to genetic variants and environmental factors. Specifically, the impact of dietary supplements and drugs on lipid remodelling must be investigated.

## 1. Introduction

The failure of clinical trials of disease modifying agents in Alzheimer’s disease (AD) highlights our limited knowledge about underlying pathophysiological mechanisms. AD often presents with diabetes co-morbidity and a wide range of metabolic perturbations occurring early in the disease process (1). APOE-ε4 is by far the strongest single gene variant contributing to increased AD risk and plays key roles in lipid transport and metabolism. Presence of the APOE-ε4 variant is correlated with higher cholesterol levels in the blood (2). More than twenty additional genes have recently been implicated in AD with functional roles in lipid processing, immune regulation and phagocytosis. Hence, both co-morbidities and known gene variants support the idea that metabolic dysfunctions may contribute to AD onset and progression.

Lipidomics methods using liquid chromatography and mass spectrometry (LC/MS) yield reliable measurements of hundreds of lipids in biological samples. LC/MS methods have been used in AD studies in attempts to define possible risk factors (3-7), diagnostic markers (8) and for highlighting novel drug targets (9-11). Perturbations in ceramides and related sphingomyelin metabolism (4, 7) were noted in many of these studies pointing towards a possible role of these lipids in aberrant signalling pathways, membrane remodelling, and apoptotic cascades during AD progression.

Changes in phosphatidylcholines were observed in several studies (11-13) pointing to a possible role for phospholipid metabolism in AD pathogenesis. Yet, AD risk prediction failed to replicate using a phosphatidylcholine (PC) biomarker panel(11, 14). Partial correlation network analysis indicated early AD biomarker Aβ_1-42_ was associated with PC and sphingomyelin (SM) (11). These studies support our hypothesis that distinct lipid biochemical pathways were dysregulated in early and late phase of AD. Here, we used LC-MS/MS based serum lipidomics analysis measured in the ADNI I cohort to define the lipid co-regulatory network of AD phenotypes, a statistical analysis tool that previously has been successfully used in the analysis of transcriptomic data (15). We investigated correlation of lipid sets with (1) Disease diagnosis, (2) CSF markers of disease Aβ_1-42_, CSF total tau and (3) cognitive decline and brain atrophy.

## 2. Material and methods

### 2.1. Study cohort

Data used in the preparation of this article were obtained from the Alzheimer’s Disease Neuroimaging Initiative (ADNI) database (adni.loni.usc.edu). The ADNI was launched in 2003 as a public-private partnership, led by Principal Investigator Michael W. Weiner, MD. The primary goal of ADNI has been to test whether serial magnetic resonance imaging (MRI), positron emission tomography (PET), other biological markers, and clinical and neuropsychological assessment can be combined to measure the progression of mild cognitive impairment (MCI) and early Alzheimer’s disease (AD). For up-to-date information, see www.adni-info.org.

The ADNI cohort information was downloaded from the ADNI data repository (http://adni.loni.ucla.edu/), supplying all the demographic information, neuropsychological and clinical assessment data, and diagnostic information that was previously published (16). Prior Institutional Review Board approval was obtained at each participating institution and written informed consent was obtained for all participants. Information about the ADNI project is provided on http://www.adni-info.org/ and the associated publication (17).

### 2.2. Pathology, clinical and lipidomics data

Untargeted lipidomics, AD diagnosis, CSF biomarkers including Total Tau (t-tau) and amyloid beta (Aβ_1-42_), cognitive decline (ADAScog13), brain atrophy represented by Spatial Pattern of Abnormality for Recognition of Early Alzheimer’s disease (SPARE-AD)(17) data were obtained from the ADNI repository (http://www.adni-info.org/). Generation and quality control of lipidomics data have been described in (18).

### 2.3. Detection of sets of co-regulated lipids

A pair-wise Spearman-rank correlation matrix was generated for lipids using the R function cor.test. The matrix was converted to a hierarchical tree model using the hclust function in R with the ward linkage method. The resulting tree model was divided into clusters using the tree cutting algorithm dynamicTreeCut (19). We used a minimum cluster size of three and a split depth of four in the tree cutting method.

### 2.4. Association modeling

Linear regression models were established for association of serum lipid abundances and CSF biomarkers and indices for cognitive decline and brain atrophy. No confounding variables were included in the regression models. Logistic regression models were calculated to associate serum lipids with AD diagnosis. Lipid abundances were scaled by the mean substracting approach. All models were unadjusted to identify all the lipid co-regulatory sets that were associated specifically with only AD or with AD and other demographics or confounding factors. Data from all ADNI-1 participants were included in the analysis.

### 2.5. Lipid set enrichment analysis

Co-regulatory lipid sets detected by the dynamicTreeCut method (19) were used as an input for cluster enrichment analysis using the Kolmogorov–Smirnov test as described in the ChemRICH method (20). In this test, the distribution of p-values was assumed to be uniform under a null hypothesis for a lipid cluster. Raw p-values obtained from the linear and logistic models were used as input for computing the enrichment statistics by comparing the experimental p-values with the uniform distribution. Set level p-values were adjusted for false discovery rate using the Benjamini–Hochberg method.

### 2.6. Source code

All statistical analyses were performed in R programming language version 3.4.0. The R-script is available at http://github.com/barupal/ADNI

## 3. Results

The main objective of the study was to identify lipid co-regulatory modules that were associated with AD diagnosis and its clinical and pathological features. In this direction, we first computed univariate association models and obtained raw p-values for each lipid. Next, we identified lipid co-regulatory modules, which were then used as set-definitions for a lipid-set enrichment analysis using the raw p-values for lipids. To minimize the false-negative rate on the set-level statistics, univariate p-values were not corrected for the multiple hypothesis testing.

### 3.1. Subject cohort and lipidomics details

Supplement Table 1 summarizes the details for the 806 ADNI participants at baseline included in the present study. The baseline ADNI1 serum lipidomics dataset contained 16 different lipid chemical classes summarizing 349 annotated lipids (Table 1).

**Table 1.** **Lipid classes and sub-classes in the ADNI serum lipidomics untargeted dataset**

| Class | Subclass | Count |
| --- | --- | --- |
| Acylcarnitine (AC) | Acylcarnitine | 9 |
| Free fatty acid (FA) | Fatty acid | 29 |
| Sterol lipids | Cholesterol | 1 |
|  | Cholesteryl ester (CE) | 8 |
| Phospholipid | Lysophosphatidylcholine (LPC) | 22 |
|  | Lysophosphatidylethanolamine (LPE) | 4 |
|  | Phosphatidylcholine (PC) | 53 |
|  | Phosphatidylethanolamine (PE) | 11 |
|  | Phosphatidylinositol (PI) | 11 |
|  | Plasmalogen phosphatidylcholine (p-PC) | 28 |
|  | Plasmalogen phosphatidylethanolamine (p-PE) | 15 |
| Sphingolipid | Ceramide (CER) | 19 |
|  | Glucosylceramide (GluCer) | 8 |
|  | Sphingomyelin (SM) | 34 |
| Acylglycerols | Diacylglycerol (DG) | 13 |
|  | Triacylglycerol (TG) | 84 |

### 3.2. Regression models for individual lipids

We first tested all individual lipids for their association with both early and late AD pathogenic markers and cognitive changes (Supplement Table S2). Raw p-values from these association models will be used as an input for the lipid set enrichment analysis in the following section. Figure 1 shows the number of significantly associated lipids in these regression models. A total of 168 lipids were found to be significant in at least one regression model (raw *p*-value < 0.05), making it difficult to biologically interpret them on individual lipid level. Two AD phenotypes showed strong positive associations with individual serum lipids, CSF Total tau and SPARE AD. Conversely, three phenotypes were mostly negatively associated with individual blood lipids, including the two related phentoypes AD diagnosis and its major contributor, ADASCog13. Overall, AD diagnosis was associated with a decline in many lipid levels which could point to lower metabolic activity in specific lipid metabolic pathways. When analyzing all individual lipids that were associated with at least one AD-phenotype, we found a very high specificity of lipid associations with a particular AD-phenotype (Figure 2 and Table S3). 48% (168/349) of all lipids were associated with at least one AD-phenotype (Table S3). Specifically, for known lipids, 63% of all AD-phenotype associated lipids were specific to only one phenotype and not shared with others (Figure 3). 28% of the detected associations of known lipids were shared between two phenotypes, driven by lipids shared between the two related phentoypes AD diagnosis and its major contributor, ADASCog13, in addition to lipids shared between total tau and SPARE-AD. Conversely, Abeta142 showed few shared lipids. Importantly, there was no identified lipid that was shared between four phenotypes, and only one lipid that was associated with all AD-phenotypes (arachidonyl-lysophosphatidylethanolamine; LPE C20:4). Many lipids are co-regulated by the activity of specific lipases or other enzymes involved in lipid remodeling. Identifying commonalities of biochemical mechanisms may lead to underlying genetic drivers or environmental factors implicated in AD-etiology. Therefore, we next focused on identifying sets of co-regulated lipids associated with AD pathophysiology rather than interpreting individual lipids.

**Figure 1.**
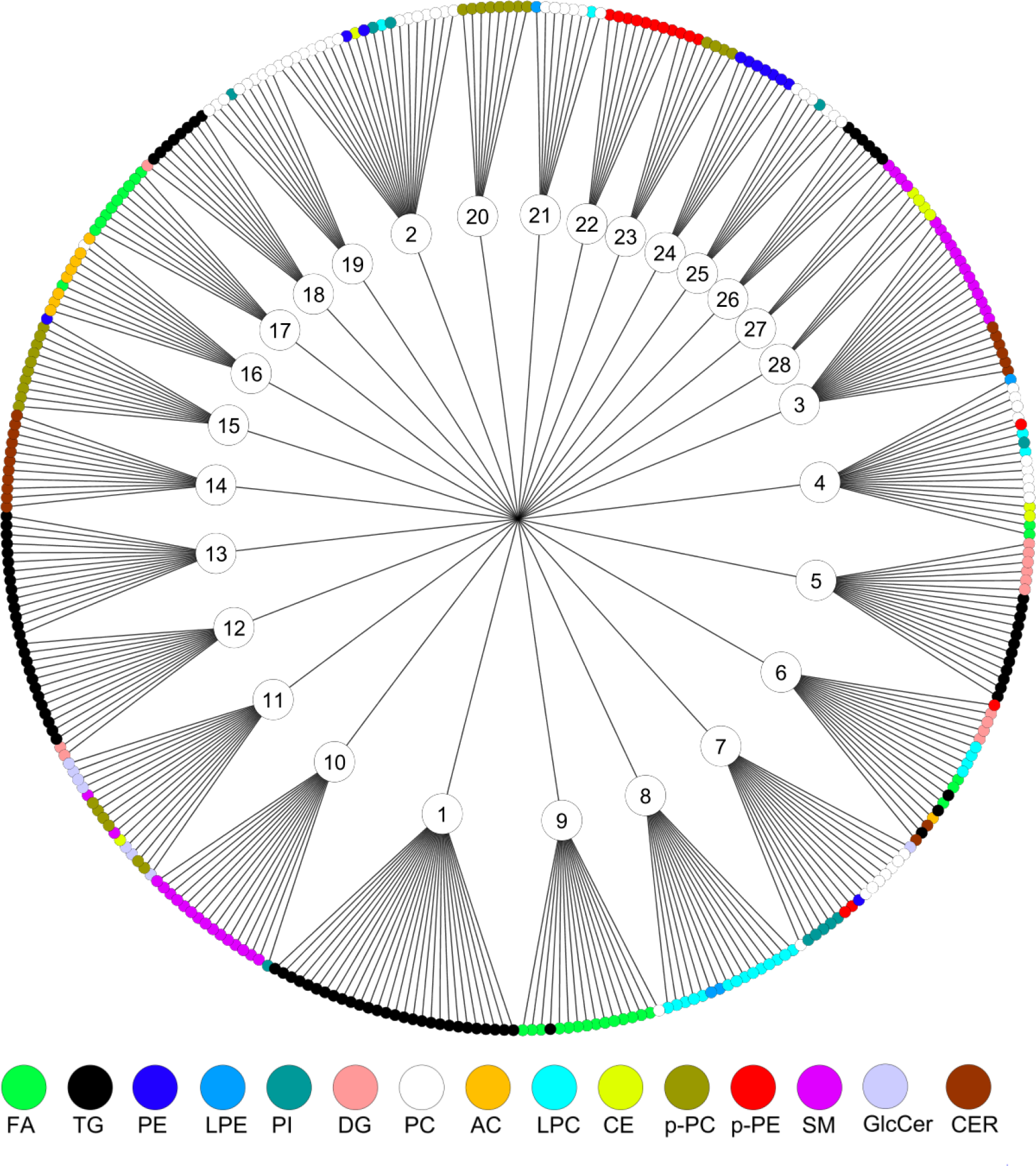
Co-regulated sets of serum lipids in the ADNI lipidomics dataset. Sets were detected by the dynamicTreeCut algorithm (see method). Node colors show different chemical classes. FA – fatty acids, AC – acyl carnitines, PC phosphatidylcholines, CE – cholesteryl esters, SM – sphingomyelins, Cer – ceramides, PE – phosphatidylethanolamines, TG – triacylglycerols, PI – phosphatidylinositols, DG – diacylglycerols, LPC – lysophosphatidylcholines.

**Figure 2.**
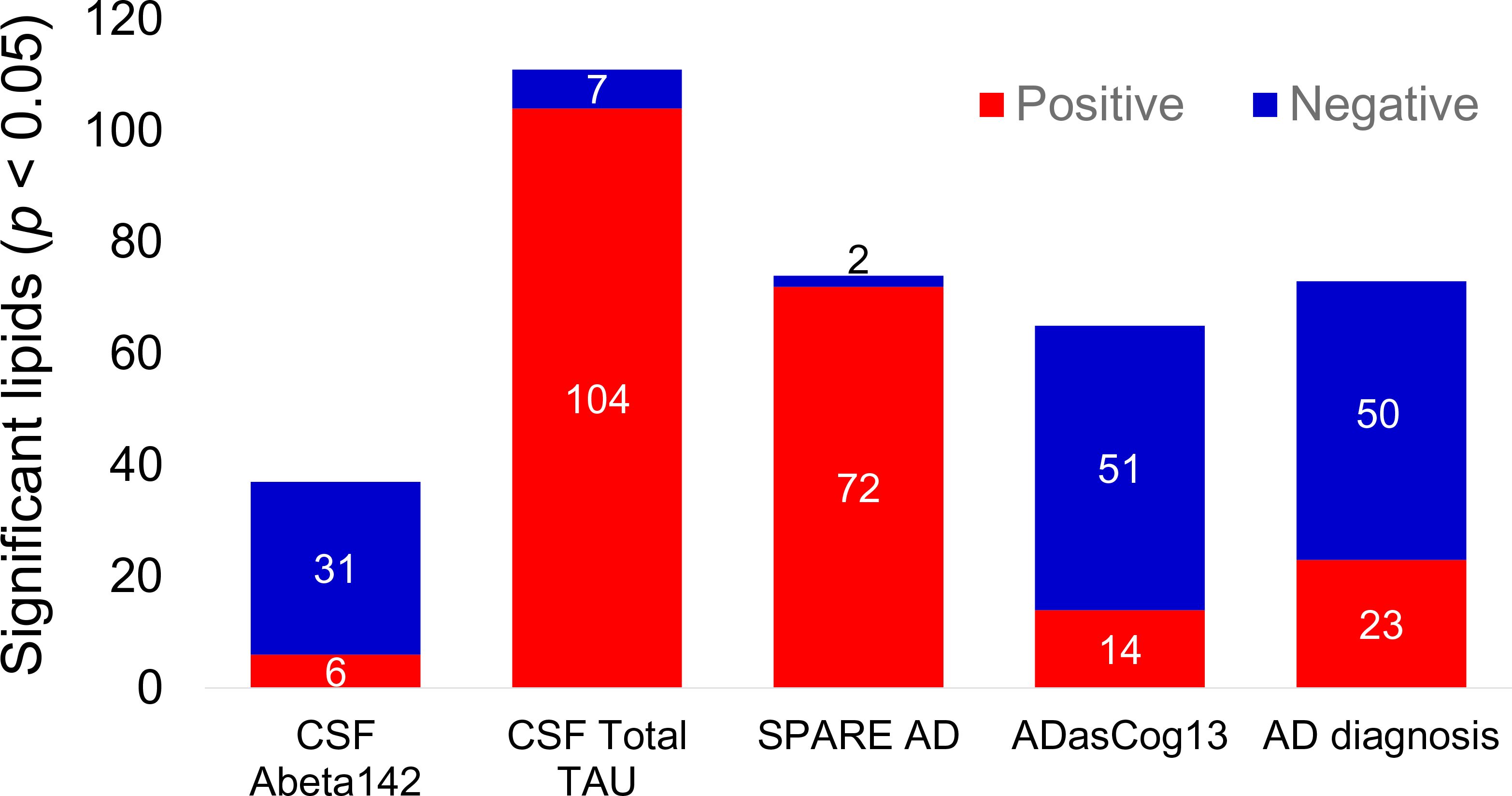
Number of significantly different lipids with AD phenotypes in univariate statistics. Directions of beta coefficients in regression models are given by colors as blue (negative) and red (positive) associations using uncorrected *p* < 0.05 values. CN : cognitively normal, LMCI : late mild cognitive impairment, AD : Alzheimer’s disease. DIAG : linear models for diagnosis, tau – linear model for tau, Aβ_1-42_ – linear model for amyloid beta peptide 42, SPARE-AD – linear model for Spatial Pattern of Abnormality for Recognition of Early Alzheimer’s disease index, ADASCog13 – Cognitive Subscale of the Alzheimer’s Disease Assessment Scale index.

**Figure 3.**
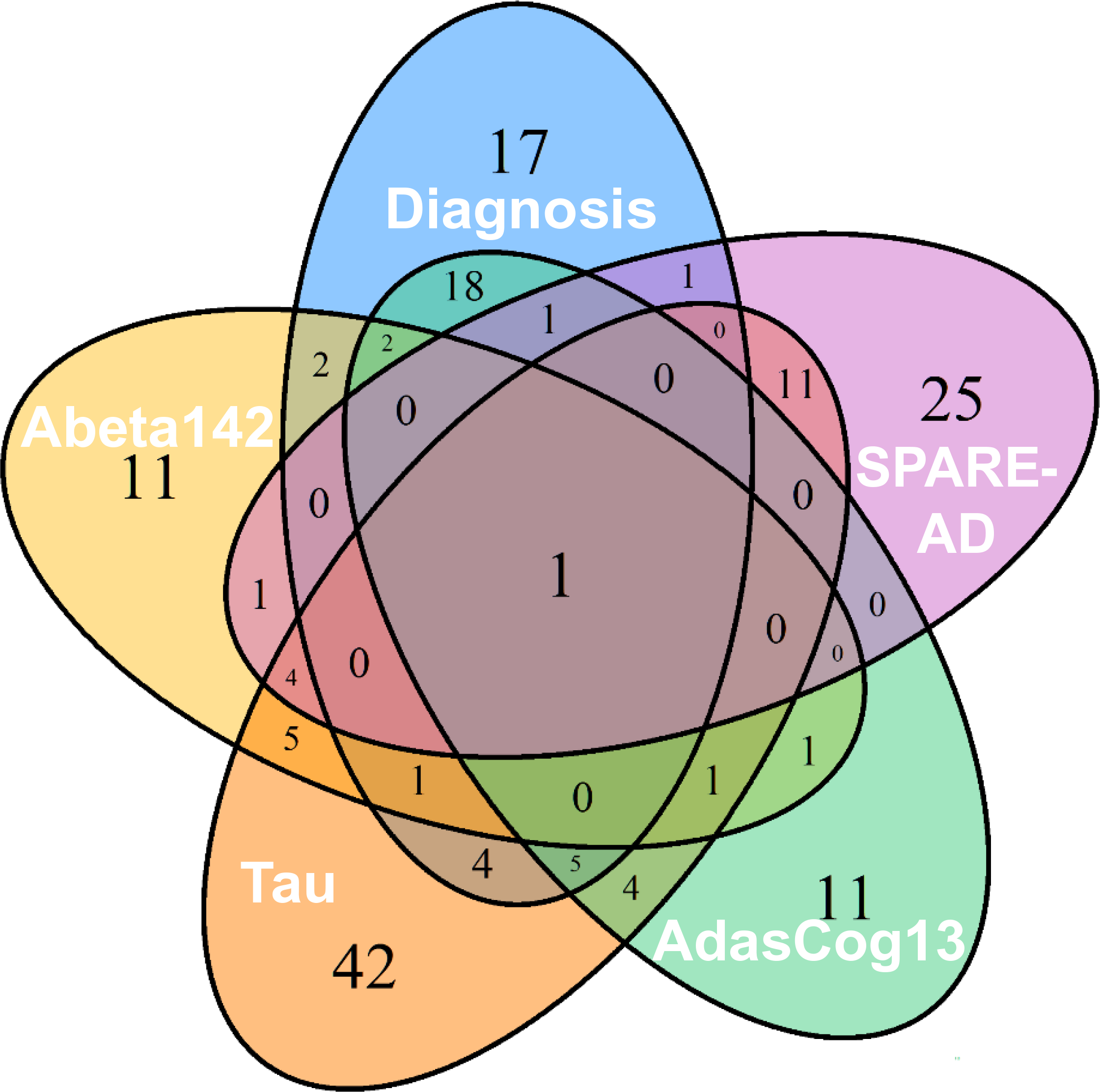
Number of significantly associated lipids across AD-phenotypes. Uncorrected *p*<0.05 values for five AD-phenotypes. DIAG : linear models for diagnosis, tau – linear model for tau, ABETA142 – linear model for amyloid beta peptide 42, SPARE-AD – linear model for Spatial Pattern of Abnormality for Recognition of Early Alzheimer’s disease index, ADASCog13 – Cognitive Subscale of the Alzheimer’s Disease Assessment Scale index.

### 3.3. Identifying sets of co-regulated lipids

Lipid classification often uses chemical structure as the only criterion. To specify the biochemical relationships between circulating blood lipids, we instead used correlation analysis to determine sets of lipids. A pair-wise Spearman correlation matrix followed by hierarchical clustering with the DynamicTreeCut dendrogram cutting method (19) yielded a total of 28 co-regulated lipid sets in the ADNI1 dataset (Figure 3). The mean size was 12.5 lipids per set, ranging from 4 to 28 members. These lipid sets (LM) were named LM1 to LM28. The average Spearman correlation coefficient *rho* across sets was 0.63 with a range of 0.19 < *rho* < 0.82. Figure 3 and Supplemental Table S2 show that some lipid co-regulatory sets were composed of lipids from the same chemical classes (such as Set-17 for free fatty acids, Set-1 for triacylglycerides and Set-14 for ceramides) whereas other sets were heterogeneous (such as Set-3 consisting of ceramides and sphingomyelins, or Set-7 that includes phosphatidylinositols and phosphatidylcholines). Interestingly, several classes of lipids were found with distinct co-regulation within each class. For example, triacylglycerides were not found as one large group of co-regulated species, but clustered in three specific sets, and similarly, free fatty acids were found in two different sets, Set-9 consisting only of saturated fatty acids and Set-17 comprised only of unsaturated fatty acids. Similarly, other lipid classes were distributed across different sets, too. For example, phosphatidylcholines were found in five sets and sphingomyelins were co-regulated in three sets, indicating downstream regulation of lipid biochemistry by specific elongases, desaturases, lipases, acyl-transferases within each lipid class (Figure 2).

### 3.4. Associating lipid sets with AD-phenotypes

These lipid groups served as input for a lipid-set enrichment analysis (LSEA)(20) along with the p-value and beta coefficient results from the regression models. Overall, 19 out of 28 lipid sets were significantly associated with at least one AD-phenotype (Figure 4, Table S3) using the Kolmogorov-Smirnoff statistical test with FDR-corrections. Eight sets were uniquely associated with only one specific AD-phenotype, but only one set was associated with four phenotypes, Set-11, consisting primarily of ceramides and phosphatidylcholines. Set-11 did not include polyunsaturated acyl chains with three or more double bonds. Six sets were associated with two AD-phenotypes and four sets were correlated with three AD-phenotypes, but no set correlated with all five phenotypes. More than two-thirds of all associations were positively correlated between lipid sets and phenotypes, mostly driven by the t-tau phenotype that also had the highest number of correlated lipid sets. Conversely, ADASCog13 showed the highest number of negative associations with lipid sets. We therefore investigated the individual phenotypes with respect to the composition of their associated lipid sets.

**Figure 4.**
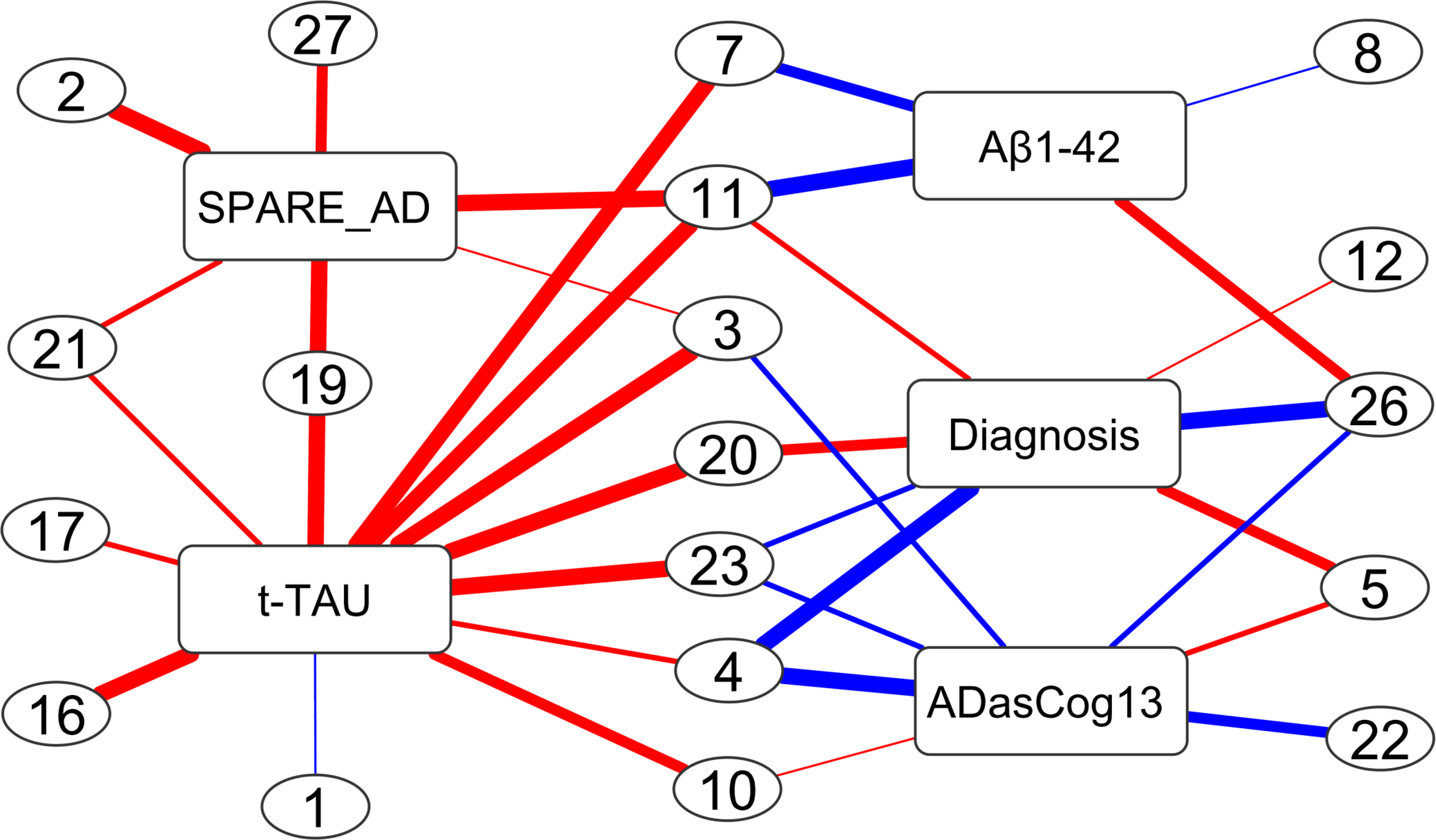
Co-regulated sets of lipids significantly associated with AD- phenotypes. Direction of associations is given by red (positive) and blue (negative) edge colors. Line thickness indicates the significance of associations (see Supplementary Table 4 for details). Lipid compositions for each set are shown in Figure 1 and Supplementary Table S1. DIAG : linear models for diagnosis, tau – linear model for tau, ABETA142 – linear model for amyloid beta peptide 42, SPARE-AD – linear model for Spatial Pattern of Abnormality for Recognition of Early Alzheimer’s disease index, ADASCog13 – Cognitive Subscale of the Alzheimer’s Disease Assessment Scale index.

#### 3.4.1. Lipid sets associated with AD diagnosis

AD-diagnosis was significantly associated with seven distinct lipid sets (Figure 4) after FDR correction. Specifically, the phenotype was highly significantly negatively correlated with lipid Set-26 and Set-4. Both sets were comprised of acyl chains with at least one polyunsaturated fatty acyl chain (PUFA) (see Table S2), either eicosapentaenoic acid (EPA), docosahexaenoic acid (DHA) or arachidonic acid (AA). Set-26 consisted exclusively of triacylglycerides that also contained either EPA or DHA, but not a single saturated fatty acid. Set-4 was mixed between different phospholipid head groups, cholesteryl esters and free fatty acids, indicating that the co-regulation mechanisms focused on the modulation and incorporation of acyl chains irrespective of the lipid class. Set-23 was also negatively correlated with AD-diagnosis and comprised of DHA-containing choline- and ethanolamine-plasmalogens. Conversely, two other sets of lipids were positively associated with AD-diagnosis, most significantly for Set-5 and Set-20, and less significantly with Set-11 and Set-12. Set-5 contained co-regulated di- and triacylglycerols with acyl groups that did not contain any PUFA with four or more double bonds, and only one single lipid with linolenic acid (C18:3). Set-20 was exclusively composed of unsaturated choline-plasmalogens, but not containing any EPA or DHA acyl chain.

#### 3.4.2. Lipid sets associated with CSF Aβ1-42

CSF Aβ_1-42_ was significantly associated with four lipid sets (Figure 4). Three sets were negatively correlated, Set-11, Set-7 and Set-8. Set-7 was the only lipid set that contained phosphatidylinositols, in addition to phosphatidyl cholines. Acyl chains were primarily saturated or mono- and di-unsaturated. Similarly, Set-8, consisted mostly of desaturated acyl groups with less than four double bonds, exclusively found as lyso-phosphatidylcholines. In the same way, no PUFA-acyl chains were found in Set-11, a heterogenous set of ceramides and choline-plasmalogens. Importantly, the only positive association of a lipid set with CSF Aβ_1-42_ was Set-26 that was completely made of PUFA- triacylglycerides.

#### 3.4.3. Lipid sets associated with CSF tau

CSF total tau correlated with 12 lipid sets, the highest number of associated lipid sets among all phenotypes (Figure 4). All sets except Set-1 were positively correlated with CSF-total tau. Three unique sets that were not shared with other phenotypes. Set-16 was composed of acylcarnitines with increasing degree of desaturation, and Set-17 was a set of monounsaturated fatty acids. Set-1 was less significant in comparison to other set associations. Four sets were shared with brain atrophy, four sets were shared with AD-diagnosis, two sets with amyloid beta and four lipid sets were shared with cognitive decline. Notably, set-19 was also associated with brain atrophy and contained mostly EPA-side chain phosphatidylcholines. Set-3 was composed of sphingomyelins and ceramides that were not associated with diagnosis or amyloid beta, but instead was also linked with cognitive decline and SPARE-AD.

#### 3.4.4. Lipid sets associated with Brain atrophy (SPARE-AD)

Brain atrophy was most significantly associated with Set 27, 19, 11 and 2 (Figure 4). Three sets (Set 2, Set 11, Set 27) were void of PUFA-side chains with either phosphoplipid or sphingolipid head groups. Conversely, Set 19 contained mostly EPA- side chain phosphatidylcholines and was further associated with CSF total tau. Similarly, Set 21 was associated with both phenotypes, containing phospholipids and their lyso-forms with the PUFA acyl chain arachidonic acid.

#### 3.4.5. Lipid sets associated with cognitive functions

Most of the lipid sets associated with cognitive decline were also associated with AD diagnosis (Figure 4). Additionally, it was negatively associated with set Set-22 which consisted of ethanolamine-plasmalogens.

## 4. Discussion

We here focused on associations between blood lipids and five AD-phenotypes guided by known contributions of lipids and metabolic co-morbidities to Alzheimer’s disease. We systematically tested both the association of individual lipids and the association with sets of co-regulated lipids. This approach showed an important advantages over previous “feature” based lipidomics-AD studies (9), (21) that did not focus on specific lipid groups, their side chains or their biological regulation. Without clear lipid identification, feature-based associations miss biological insights and have a high risk of not being validated in subsequent studies because each individual lipid may cause more than one feature in LC-MS based lipidomics(21, 22). Instead we used the largest published AD-lipidomics data set (18) to date with 349 identified lipids belonging to 13 major lipid classes, identified by extensive MS/MS fragmentation analysis (23) and enabling analyses reaching to the level of acyl chains. A second difference to previous efforts was a focus on summarizing lipids by statistical co-regulation instead of only relying on univariate analysis or grouping by lipid head groups. This expansion of classic statistical analysis was critical to extend from diagnostic biomarkers (that need correctiong for multiple testing using false discovery rate (FDR) adjustments) to revealing underlying biological mechanisms. The axiom of univariate analyses, the mutual independence of variables, is untrue in lipid biology. Moreover, stringent FDR corrections lead to an increased number of false negative results and compromise the statistical power to investigate the biological mechanisms and pathways. As lipids are poorly presented in biochemical pathway databases (20), classic metabolic pathway enrichment analysis (24) ignores a majority of detected lipids and is unsuitable for lipidomics. Instead, Spearman-rank correlation based matrices yielded specific clusters of lipids associated with Alzheimer’s disease phenotypes by using the robust Kolmogorov-Smirnov test for *p*-value distributions. These lipid sets showed very distinct metabolic features that we identified as preferential use of specific polyunsaturated fatty acids that replaced saturated or monounsaturated fatty acids for distinct lipid classes. These mechanisms of lipid desaturation, elongation, and acyl-chain remodeling were disturbed in early and later stages of Alzheimer disease. A minimal overlap among lipid sets was observed (Figure 3) with respect to statistical associations with AD phenotypes, indicating that quite distinct lipid biochemical processes were involved in the early and later stages of AD. Lipid metabolic pathways associated with the early stage, in particular, may provide new therapeutic targets to stop AD progression. MUFA-containing lipids were positively associated with the brain atrophy and tau accumulation whereas PUFA-containing lipids were negatively associated with AD diagnosis and cognitive decline. Therapeutic strategies targeting MUFA lipid pathways at the early stages of AD could therefore be potentially more effective in delaying the progression of the disease.

### 4.1 Lipids linked to the amyloid beta clearance pathway

A decrease in the **CSF Aβ_1-42_** peptide marker is indicating a poor clearance of the peptide in the brain, leading to its extra-neuronal accumulation. In our study, poor clearance was indicated by negative associations with lipids sets, including sets that contained phosphatidylinositols, lysoPCs, ceramides and choline-plasmalogens and PUFA TGs. The amyloid β peptide is known to cause mitochondrial dysfunction (25) which can lead to neurodegeneration via autophagic cascades (26, 27). The associated lipids, specifically ceramides and phosphatidylinositols and lysoPCs have been linked with cell death and may also contribute in the Amyloid Beta mediated toxicity in neurons (28-30). Higher levels of ceramides containing oleic acid (C18:1) have been shown to increase AD risk (4, 5). We validate this finding in our study and also observed lower levels of phosphoinositols containing polyunsaturated fatty acids to correlate with poor Amyloid-beta clearance. An alternative explanation to our data is an impaired amyloid beta clearance in the liver(31)that subsequently leads to dysregulation of lipid metabolism in the liver. Overall, our data suggest that these lipid sets can serve as serum biomarkers for disturbed Amyloid beta pathway regulation in brain and can complement Amyloid beta imaging assays.

### 4.2 Cerebrospinal fluid total tau

CSF tau is a marker for accumulating tau tangles in the brain tissues, causing neurodegeneration. We found that the total CSF tau marker was significantly associated with lipid sets enriched in monounsaturated fatty acids, acyl-carnitines, ceramides, sphingomyelins, and EPA containing phosphatidylcholines. Increased fatty acids and acyl-carnitines are known markers of impaired fatty acid beta oxidation in mitochondria(32), specially during metabolic diseases such as diabetes and obesity(33, 34). We found free fatty acids and acylcarnitines to be positively correlated with total tau, supporting the notion of tau mediated neurodegeneration and mitochondrial dysfunctions. Mitochondrial impairment was further evidenced by positive associations of total tau with sets of ceramides, because accumulating ceramide are known to induce cell death and to increase the AD risk in normal subjects (5) Rozen et al. 2011. Higher ceramide levels were also reported for early stage Alzheimer’s disease (35-37). These findings indicate that these lipids may be involved in early neurodegenerative pathways, and their underlying pathways might lead to candidates for new therapeutic strategies.

### 4.3 Lipid sets linked with brain atrophy

SPARE-AD is a composite index of brain atrophy and indicates the neurodegeneration magnitude. We found a high overlap of lipid sets that were associated with both SPARE-AD and total tau, reinforcing the usability of these serum lipids as biomarkers for neurodegeneration. These lipid sets included phosphatidylcholines and sphingolipids that were enriched in polyunsaturated fatty acyls (PUFA) eicosapentaenoic acid and arachidonic acid (EPA, AA). These fatty acids are main components of brain lipids(38, 39). The loss of brain tissue may cause an increase in levels of serum lipids that include EPA and AA as acyl groups through lipid remodelling (40, 41). We here identify lipid pathways associated with tau-mediated brain atrophy that eventually contributes to AD.

### 4.4 AD diagnosis and cognitive decline

Previous publications reported that lower levels of PUFA in AD subjects across multiple lipid classes, along with higher levels of monounsaturated lipids (4, 8, 9, 42-46). We found numerous, very significant associations of omega-3 and omega-6 containing complex lipids with AD diagnosis and cognitive functions. Our analysis is consistent with these results as shown by AD associated lipids in Set-4, Set-20, Set-23 (Figure 4). We here specify that the major difference is not related to total levels of mono- or polyunsaturated fatty acids, but the extent at which these fatty acids are incorporated into different complex lipids. Clear differences in circulating PUFA phospholipid and PUFA triacylglycerol levels in AD subjects in comparison to normal subjects were observed, likely due to dysregulation of biosynthesis in liver. Here, substrate preference of MGAT and DGAT enzymes in the liver may play an important role, but the exact specificities of acyltransferase enzymes (and their corresponding lipase enzymes) are not well studied. Levels of anti-inflammatory plasmalogens (47), important lipids for brain membrane functions (45, 48), were decreased in AD patients in comparison to cognitively normal subjects. Lower levels of plasmalogens have been linked to AD (45). However, in clinical trials, EPA and DHA supplementation do not improve the cognitive function of AD subjects (49). Nutritional intervention trials such as the European LipiDiDiet have failed to show any cognitive improvement in AD subjects. A broad-spectrum effect of FOS on additional lipid pathways may explain the failure of this trial and warrants further lipidomics studies for serum specimens of this trial’s participants. It was observed that the incorporation of omega-6 fatty acids was increased in AD subjects. These fatty acids are well-known precursors to pro-inflammatory molecules such as leukotrienes. Further studies are needed to test if post-mortem brain tissues of AD subjects show similar disruption in fatty acid incorporation and if these patterns correlate with AD severity or other AD phenotypes.

## 5. Conclusions

Using co-regulated sets of lipids enabled us to find significant associations of lipids with Alzheimer’s disease that led to biochemical mechanisms. Across the spectrum of AD progression, pathways were dysregulated that pointed to lipid desaturation, elongation and remodelling. These pathways provide new targets as well candidate biomarkers for the population screening for AD prevention. Future studies are needed to tease out the roles of genetic variations, drug, and diet the metabolism of MUFA and PUFAs and their complex lipids and their roles in AD.

## Acknowledgments

**Funding**

This work was funded through NIH awards U54 AI138370, U19 AG023122 and U2C ES030158. National Institute on Aging (R01AG046171, RF1AG051550, and RF1AG057452 and 3U01AG024904-09S4) supported the Alzheimer Disease Metabolomics Consortium which is a part of NIA national initiatives AMP-AD and M2OVE AD. Data collection and sharing for this project was funded by the Alzheimer’s Disease Neuroimaging Initiative (ADNI) (National Institutes of Health Grant U01 AG024904) and DOD ADNI (Department of Defense award number W81XWH-12-2-0012). ADNI is funded by the National Institute on Aging, the National Institute of Biomedical Imaging and Bioengineering, and through generous contributions from the following: AbbVie, Alzheimer’s Association; Alzheimer’s Drug Discovery Foundation; Araclon Biotech; BioClinica, Inc.; Biogen; Bristol-Myers Squibb Company; CereSpir, Inc.; Cogstate; Eisai Inc.; Elan Pharmaceuticals, Inc.; Eli Lilly and Company; EuroImmun; F. Hoffmann-La Roche Ltd and its affiliated company Genentech, Inc.; Fujirebio; GE Healthcare; IXICO Ltd.; Janssen Alzheimer Immunotherapy Research & Development, LLC.; Johnson & Johnson Pharmaceutical Research & Development LLC.; Lumosity; Lundbeck; Merck & Co., Inc.; Meso Scale Diagnostics, LLC.; NeuroRx Research; Neurotrack Technologies; Novartis Pharmaceuticals Corporation; Pfizer Inc.; Piramal Imaging; Servier; Takeda Pharmaceutical Company; and Transition Therapeutics. The Canadian Institutes of Health Research is providing funds to support ADNI clinical sites in Canada. Private sector contributions are facilitated by the Foundation for the National Institutes of Health (www.fnih.org). The grantee organization is the Northern California Institute for Research and Education, and the study is coordinated by the Alzheimer’s Therapeutic Research Institute at the University of Southern California. ADNI data are disseminated by the Laboratory for Neuro Imaging at the University of Southern California.

## Author’s contribution

DKB and OF generated the lipidomics dataset. DKB, RB, RKD and OF design the study. DKB and SF performed the statistical analyses. AJS, PJM, MA and KN contributed in data interpretation. All author contributed in the manuscript writing.

Transparency declaration: Authors affirms that the manuscript is an honest, accurate, and transparent account of the study being reported; no important aspects of the study have been omitted; and any discrepancies from the study as planned have been explained.

## ABBREVIATIONS

FA: fatty acids
AC: acetyl carnitine
PC: phosphatidylcholine
CE: cholesteryl esters
SM: sphingomyelins
Cer: Ceramide
PE: phosphatidylethanolamine
TG: triacylglycerol
PI: phosphatidylinositol
DG: diacylglycerol
LPC: lysophosphatidylcholine
Aβ: Amyloid Beta
SPARE-AD: Spatial Pattern of Abnormality for Recognition of Early Alzheimer’s disease
LCRS: Lipid Co-Regulatory Set
MUFA: mono unsaturated fatty acid
PUFA: poly unsaturated fatty acid
ADASCog13: Alzheimer’s Disease Assessment Scale–Cognitive subscale
ADNI: Alzheimer’s disease neuroimaging initiative
EPA: eicosapentaenoic acid
DHA: docosahexaenoic acid
CSF: cerebrospinal fluid

